# Complex associations of environmental factors may explain Blanchard’s Cricket Frog, *Acris blanchardi* declines and drive population recovery

**DOI:** 10.1101/272161

**Authors:** Malcolm L. McCallum, Stanley E. Trauth

**Affiliations:** P.O. Box 150, Langston, Oklahoma 64040.; Aquatic Resources Center, School of Agriculture & Applied Sciences, E.L. Holloway Agriculture Research, Education, & Extension, Langston University, P.O. Box 1730, Langston, Oklahoma 73050, USA; P.O. Box 599, Department of Biological Sciences, Arkansas State University, State University, Arkansas 72467.

## Abstract

Blanchard’s Cricket Frog, *Acris blanchardi*, is a small hylid frog that was once among the most common amphibians in any part of its range. Today, it remains abundant in much of the southern portion of its range, but is now disappearing elsewhere. Our analysis of habitat characters observed across several states revealed interesting relationships of these factors with the abundance or presence of Blanchard’s Cricket Frog. Further, we later established two ½ acre ponds based on these relationships that led to immediate colonization of the ponds by cricket frogs followed by explosive production of juveniles less than a year later. Our findings suggest that habitat management for this species should specifically manage the shoreline grade and especially the aquatic floating vegetation to maximize population growth and sustenance.

## Introduction

Conservation strategies are impaired when we lack a broad understanding of target’s natural history (Bury 2006; McCallum and McCallum 2006; Gilpin 1986). In fact, the progress of an R or D extinction vortex pivots on the expression of key life history traits such as reproduction potential and offspring survivorship (Gilpin 1986). An R vortex is one in which a disturbance facilitates reduction in population size, thus increasing vulnerability to additional disturbances. Herein, we hypothesize that floating mats of vegetation may factor into whether populations of Blanchard’s Cricket Frog (*Acris blanchardi*) remain robust, or follow the path of an R extinction vortex to extirpation.

Blanchard’s Cricket Frog is a small, short-lived frog (Burkett 1984; McCallum et al. 2011; Letinhen et al. 2011) previously designated as a subspecies of the Northern Cricket Frog (*A. crepitans*) (McCallum and Trauth 2006; Gamble et al. 2008) and currently (but questionably) encompassing the Coastal Cricket Frog (*A. blanchardi paludicola* [Rose et al. 2006]). It is known for its color polymorphism (Pyburn 1958; 1961; Gray 1972; 1983) and zig zag jumping patterns (McCallum 2011).

The color polymorphism refers only to the color or presence of a dorsal stripe running down its back (Gorman 1986; Gray 1984; Milstead et al. 1974), which results from multiple alleles at one locus (Pyburn 1961). However, color diversity in this species is remarkable with spots and splotches of assorted sizes ranging from green to red to brown, and combinations thereof (McCallum Pers. Obs.; Gray and Pantex 1995; Cope 1889). Some of the color variation may be seasonal metachrosis (Cope 1889; Gray 1972, 1983; Strate 1987). There is evidence that color morph frequency is related to variation in the stream substrate color or due to the presence of vegetation (Nevo 1973a; Pyburn 1961). However, laboratory results using frogs housed in aquaria with American Bullfrogs (*Lithobates catesbeianus*) or Common Garter Snakes (*Thamnophis sirtalis*) did not support this hypothesis (Gray 1978; 1984).

When a cricket frog flees from a predator, it frequently jumps 2-3 times at relatively consistent angles away from the predator’s attack to optimize distance and angular displacement from the lunging opponent and then scurries into a terrestrial or aquatic abode (McCallum 2011). Terrestrially, it will bury itself into a small grassy patch. If it jumps in the water it will often swim back to hide in shoreline vegetation (Gray 1978; McCallum pers. obs.). If it dives, it will often enter a patch of submerged vegetation or floating filamentous algae and remain motionless, floating with the vegetation and the current (McCallum 1999; 2011). Alternatively, it will dive to the bottom, into the mud, remain motionless, and allow the mud to settle on it back. If leaves or other loose materials are on the bottom, it will hide under them. It often avoids moving after being poked or touched while hiding in a patch of vegetation, as if dead (McCallum 1999). It also death-feigns occasionally when held (McCallum 1999). Then, suddenly it jumps away or swims zig-zagged along the bottom and dives again into the benthos or a different aquatic vegetation patch.

These frogs overwinter in cracks and burrows in the soil (Gray 1971; Badje et al. 2016), gravel along streambeds (McCallum and Trauth 2003a) and other terrestrial locations (Irwin et al. 1999; Badje et al. 2016; Kenney et al. 2012; Swanson and Burdick 2010). Outside of hibernation, they tend to occur in terrestrial settings (Smith et al. 2003) within 5 – 6 cm of a vegetation patch and 10 – 50 cm from shore (Burdick and Swanson 2010). However, they sometimes disperse considerable terrestrial distances between wetlands (Gray 1983; McCallum 2003; McCallum et al. 2011; McCallum et al. 2003; Youngquist and Boone 2014). Breeding takes place March – October, depending on the region, and multiple clutching is prominent outside of South Dakota (McCallum et al. 2011; McCallum and Trauth 2004). Males call both during day and night (McCallum and McCallum 2018). Young-of-the-year males are capable of reproduction within months of metamorphosis, but females remain immature until the following summer (McCallum et al. 2011). Through most of its range these frogs live one year and the population turns over by October (Lehtinen and McDonald 2011; McCallum et al. 2011). However, in more northerly portions of its range (e.g. New York, South Dakota) they may persist a second year (McCallum and Trauth 2004; 2011).

Blanchard’s Cricket Frog is declining in the Northern parts of its range and it has conservation status in New York (Kenney and Stearns 2015), Michigan (Michigan Natural Features Inventory 2007; Lehtinen 2002), Minnesota (Minnesota Department of Natural Resources 1996), Wisconsin (Wisconsin Department of Natural Resources 2015), South Dakota (Fischer et al. 1999), and appears extirpated from Colorado (Hammerson and Livo 1999); West Virginia (Dickson 2002) and Ontario (Canada; Beauclerc et al. 2010). Reasons for the declines may relate to climate change (McCallum 2010; Morgan 2016), contaminants (Reeder et al. 1998; 2005; Russel et al. 2002; Hayes et al. 2011), habitat acidification (Lehtinen and Skinner 2006), wetland management (Lannoo 1998; Gordon et al. 2016; Krynak et al. 2016), and other yet unclarified reasons (Burdick a 2016; McCallum and Trauth 2003b; Dickson 2002). Chytridiomycosis (Steiner and Lehtinen 2008; Zippel and Tabaka 2008), lymphedema (McCallum 2018) and anatomical abnormalities (McCallum and Trauth 2003b; 2004; Gray 2000a; 2000b) have been reported, sometimes preceding extirpation.

The importance of aquatic vegetation for this species has been hypothesized (Nevo 1973a; Regan 1973; Pyburn 1961; McCallum 2003; but see Gray 1978; 1984). Herein, we combine numerous field observations with a structured, short-term study to understand the importance of floating vegetation mats and other habitat features to Blanchard’s Cricket Frog.

## Materials and Methods

The structured study was conducted at the Mammoth Spring National Fish Hatchery (MSNFH) in Mammoth Spring, Arkansas. Additional observations were made in Illinois, Kansas, Louisiana, Michigan, Missouri, Oklahoma, and Texas (Appendix 1). The MSNFH has 15 culture ponds (Fig. 1) used for raising aquatic organisms. The ponds are approximately 10 m apart. The staff of the MSNFH monitored water quality and maintained similar ranges of standard water quality parameters in all of the ponds. We visited MSNFH during June – August 2000. On each visit, we estimated the percent of the water surface covered by floating mats of Duckweed (*Lemna major* and *Lemna minor*]) interspersed with filamentous algae (*Spirogyra sp*. and *Clodophora sp*.). Then, the perimeter of each pond was surveyed between 1100 and 1300 hrs (military time, this is the peak activity period for Blanchard’s Cricket Frog [pers. observ.]). Blanchard’s Cricket Frogs were tallied at each pond, noting general habitat parameters (Appendix 1). Minitab 13.3 was used for statistical analysis. Abundance data was tested for normality using the Anderson-Darling normality test, and non-normally distributed data was transformed using the normalize function in Minitab 13.0. Regression analysis used each individual pond and percent vegetation coverage as predictors of frog abundance. An α = 0.05 was applied for decision theory. Various isolated observations recorded from 1979 to present from sites among numerous Midwestern sites within the frog’s range were incorporated into the analysis (Appendix 1). Abundance at each location was qualitatively scored on a scale from 0 – 5 for each location (0 = absent, 5 = extremely abundant). The abundance scores were statistically tested for associations with shoreline slope < 30° vs > 30°, presence of numerous general habitat structures. Definitions of vegetation/habitat were necessarily broad to encompass interpretations of notes from the author’s field journals. These predictors were classified as present (=1) or not present (=0) and subjected to regression analysis to identify relationships among habitat features. Strongly correlated variables were removed from the predictor set. Then the data were subjected to best subsets regression to identify the best predictors. The regression coefficients of the resulting models with the lowest Mallow’s C_p_ Statistic and variance were compared. The five models with the lowest variance and C_p_ were subjected to multiple linear regression. The best regression models were selected by comparing the valence inflation factor (VIF) and predicted residual error sum of squares (PRESS). Durbin-Watson statistic was used to identify potential autocorrelation. Upon analysis of these results, some predictors were combined because of the fine distinctions made between some habitat types.

**Figure 1.**
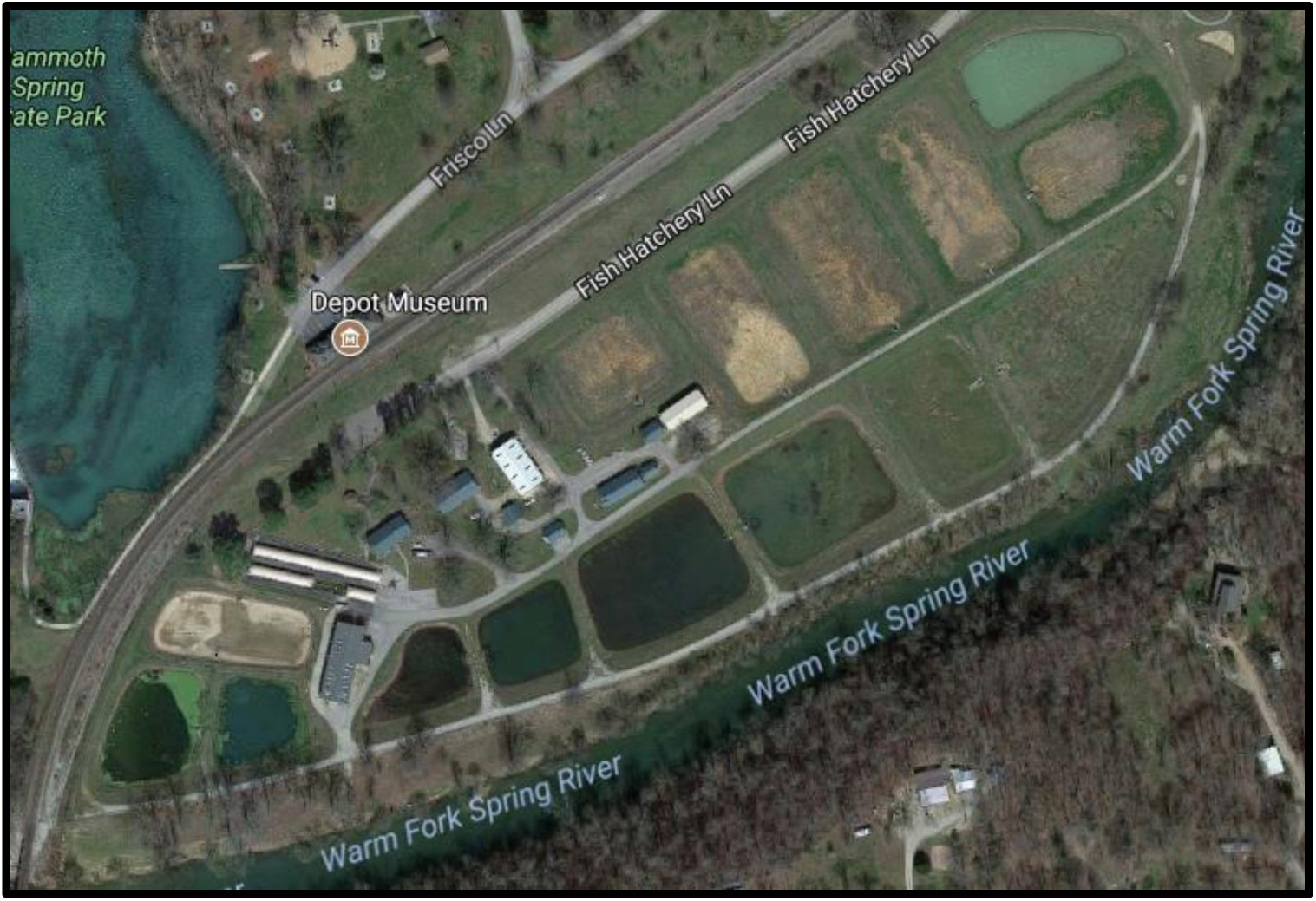
The Mammoth Spring National Fish Hatchery (Mammoth Spring, Arkansas).

Upon completing these analyses in July 2016, two 1/2 acre, 2.5 m deep artificial ponds in the vicinity of Coyle, Oklahoma were flooded with water without aeration or fish, and aquatic *Chara* sp. and floating algal mats allowed to grow unrestrained. Frog populations were counted after one and two months. Then, in July 2017 the anuran population was examined to determine if application of key habitat features would benefit Blanchard’s Cricket Frog.

## Results

### General observations

A small private pond in Collinsville, Illinois had thick mats of duckweed extending over 100% of the surface during the summer from 1979-84. Blanchard’s Cricket Frog was very abundant there. In the summer of 1984, the lake owners introduced carp to control the vegetation. The following year, both the duckweed mats and Blanchard’s Cricket Frog were absent from the lake.

Large numbers of Blanchard’s Cricket Frogs called from mats of *Myriophyllum* sp. and *Ceratophyllum sp*. April-August 2000-2002 at Jane’s Creek (Ravenden Springs, Arkansas). Late in the season (June-August) large numbers of new metamorphs used this same habitat. Upon changes in the streambed after repair of the bridge (August 2002), this mat was gone, and the reduced abundance of Blanchard’s Cricket Frogs at that location (2002-2003) dropped from 4.5 metamorphs/m^2^ (July 2000-2002) to 2 metamorphs/m^2^ in July 2003.

### Structured field study

I observed 1,178 frogs among all ponds at the Mammoth Spring Fish Hatchery, and 38 (SE = 12.5) frogs/pond/ visit. Both the pond (T = 2.87, *P* = 0.008) and percent vegetation coverage (T = 7.60, *P* < 0.001) interacted to predict the abundance of Blanchard’s Cricket Frog (*r*^2^ _2,28_ = 0.719, *P* < 0.001). In the four ponds that had < 5% vegetation and Blanchard’s Cricket Frogs present, the frogs occurred in pockets of emergent vegetation interspersed with Duckweed and algal mats (Table 1).

**Table 1.** Estimated floating vegetation coverage versus abundance of Blanchard’s Cricket Frog (*Acris blanchardi*) at Mammoth Spring National Fish Hatchery (Mammoth Spring, Arkansas). D = pond was drained, no water present.

| Pond # | 19 June | # frogs | 19 July | # frogs | 7 August | # frogs |
| --- | --- | --- | --- | --- | --- | --- |
|  | % veg |  | % veg |  | % veg |  |
| 1 | < 5 | 0 | < 5 | 0 | < 5 | 0 |
| 2 | 10 | 19 | < 5 | 4 | 25 | 89 |
| 3 | 40 | 45 | 40 | 41 | 40 | 31 |
| 4 | < 5 | 1 | < 5 | 1 | < 5 | 0 |
| 5 | < 5 | 0 | < 5 | 0 | < 5 | 0 |
| 6 | D | 1 | 50 | 101 | 50 | 56 |
| 7 | D | 0 | D | 0 | D | 0 |
| 8 | < 5 | 7 | 10 | 36 | -- | -- |
| 9 | D | 0 | 33 | 377 | -- | -- |
| 10 | 25 | 47 | 20 | 20 | -- | -- |
| 11 | 20 | 13 | < 5 | 3 | -- | -- |
| 12 | 30 | 64 | 30 | 31 | -- | -- |
| 13 | < 5 | 13 | 30 | 87 | -- | -- |
| 14 | 20 | 43 | < 5 | 8 | -- | -- |
| 15 | 20 | 41 | D | 0 | -- | -- |

### Analysis of Field Notes

The results from analysis of field notes produced five models with VIF < 3.1 and S = 1.1896 – 1.2120 and *r*^2^ = 0.573 – 0.504 (Table 2). Floating mats of sphagnum, duckweed/algae, and submerged vegetation along the water surface were positive predictors in 71.0% of the 31 models produced from best subsets regression. A shoreline with a slope > 30° was a negative predictor of cricket frog abundance in 83.9% of the 31 models produced. These factors appeared in all of the five best models predicting abundance of Blanchard’s Cricket Frog. The presence of gravel shorelines with light vegetation was a positive predictor in 58.1% of the models and gravel shorelines without vegetation was a positive predictor in 51.6% of the models. Three of the five best models included unvegetated gravel shorelines, and two included vegetated gravel shorelines. None of the five best models included both types of gravel shorelines. Multiple linear regression analysis of each of the five best models revealed that there was lack of fit for the models with greater than five predictors (Appendix 2) and that the influence of light vegetation on gravel banks was minimal.

**Table 2.** Five best regression models based on outcomes of best subsets regression.

| $r^2$ | $C_p$ | S | Predictors |
| --- | --- | --- | --- |
| 0.573 | 2.7 | 1.2120 | GBW, SFM, DW, SV, $> 30^\circ$ |
| 0.573 | 2.7 | 1.2120 | GB, SFM, DW, SV, $> 30^\circ$ |
| 0.593 | 2.4 | 1.1948 | GBW, MFV, SFM, DW, SV, $> 30^\circ$ |
| 0.593 | 2.4 | 1.1948 | GB, MFV, SFM, DW, SV, $> 30^\circ$ |
| 0.604 | 3.1 | 1.1896 | GBW, MF, MFV, SFM, DW, SV, $> 30^\circ$ |

Reanalysis of these data after combining the various vegetation mats and the gravel bank types revealed that that there was still high correlation among RGV and other predictors so it was removed from the analysis. Best subsets regression identified the five best models among 23 combinations of possible predictors for frog abundance (Table 3) with *r*^2^ = 0.603 – 0.634. The five best models had C_p_ = 1.6 – 2.1 and variances of 1.1225 – 1.1478. Thick floating mats of vegetation were predictors of frog presence in 96.0% of the models, and all five of the best relationships identified. A slope greater than 30° was an important predictor of frog absence in 87.0% of the 23 models generated and four of the five best models identified. Gravel banks predicted frog presence in 78.2% of the generated models and all of the five best models. Cattails and/or thick tall grasses on the shoreline predicted absence of frogs in 78.2% of the generated models, but were used in three of the best five models. No autocorrelation was identified; however, there was lack of fit in all of the new models except for

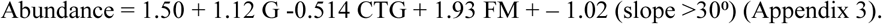

**Table 3.** Five best regression models based on outcomes of best subsets regression after combining some predictors. (G = gravel banks, FM = floating vegetation mats, SB = sand bottom, CTG = cattails or tall grasses, 30° = slope of the stream or pond bank).

| r<sup>2</sup> | Cp | S | Predictors |
| --- | --- | --- | --- |
| 0.603 | 1.7 | 1.1478 | G, FM, > 30° |
| 0.619 | 1.6 | 1.1349 | G, SB, FM, > 30° |
| 0.617 | 1.8 | 1.1368 | G, CTG, FM, > 30° |
| 0.634 | 1.6 | 1.1225 | G, CTG, SB, FM, > 30° |
| 0.630 | 2.1 | 1.1287 | G, CTG, SB, FM, < 30° |

By May 2017, the artificial ponds in Oklahoma had 25% surface coverage by algal mats and aquatic vegetation was evident. The condition of ponds by July 2017 is pictured in Figure 2. Blanchard’s Cricket Frog did not immediately colonize the two newly flooded artificial ponds. During Spring 2017, they began colonizing the ponds with calling choruses at each pond easily exceeding 50 frogs. By July 2017, the population had exploded as seen in provided video (Fig. 3 and Supplemental materials). Each splash in the water is an individual frog entering it from the shoreline. Also, of note is the immense number of juveniles utilizing the floating vegetation across the surface of each pond (Fig. 3 and supplemental materials). Finally, this population growth was accomplished despite a strong American Bullfrog population (199 and 211 bullfrogs in each pond as of May 2017).

**Figure 2.**
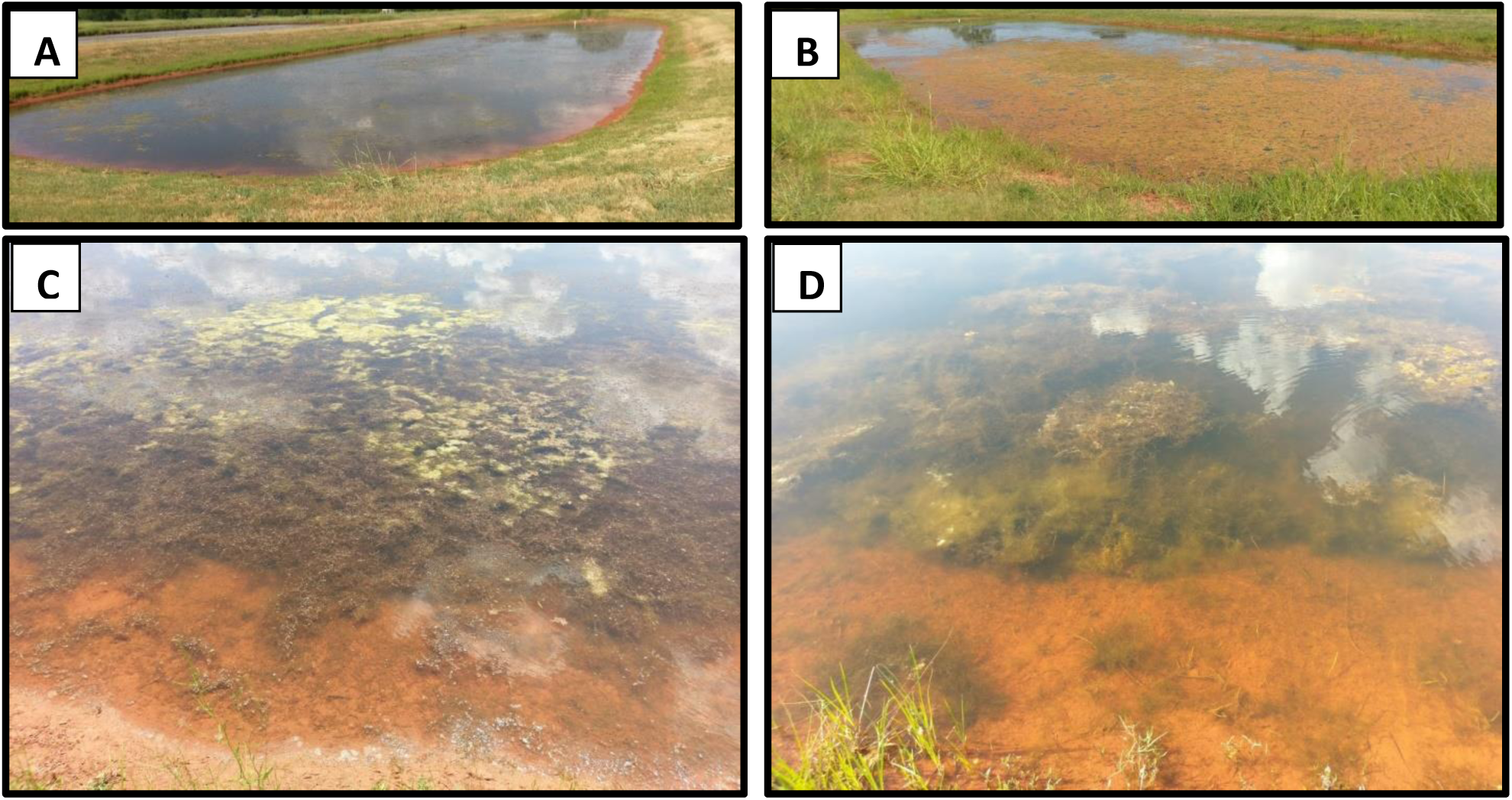
Two one-half-acre, fishless ponds with aquatic vegetation. Although the shoreline appears sloped in the photo, the angle at the water’s edge is very low grade. Grass was mowed along the edges to maintain short vegetation. Ponds are labeled A and B for reference in Fig. 3 and 4. C) *Chara* sp. interspersed with *Cladaphora* sp. in Pond A. D) Filamentous algae dominating aquatic vegetation in Pond B.

**Figure 3.**
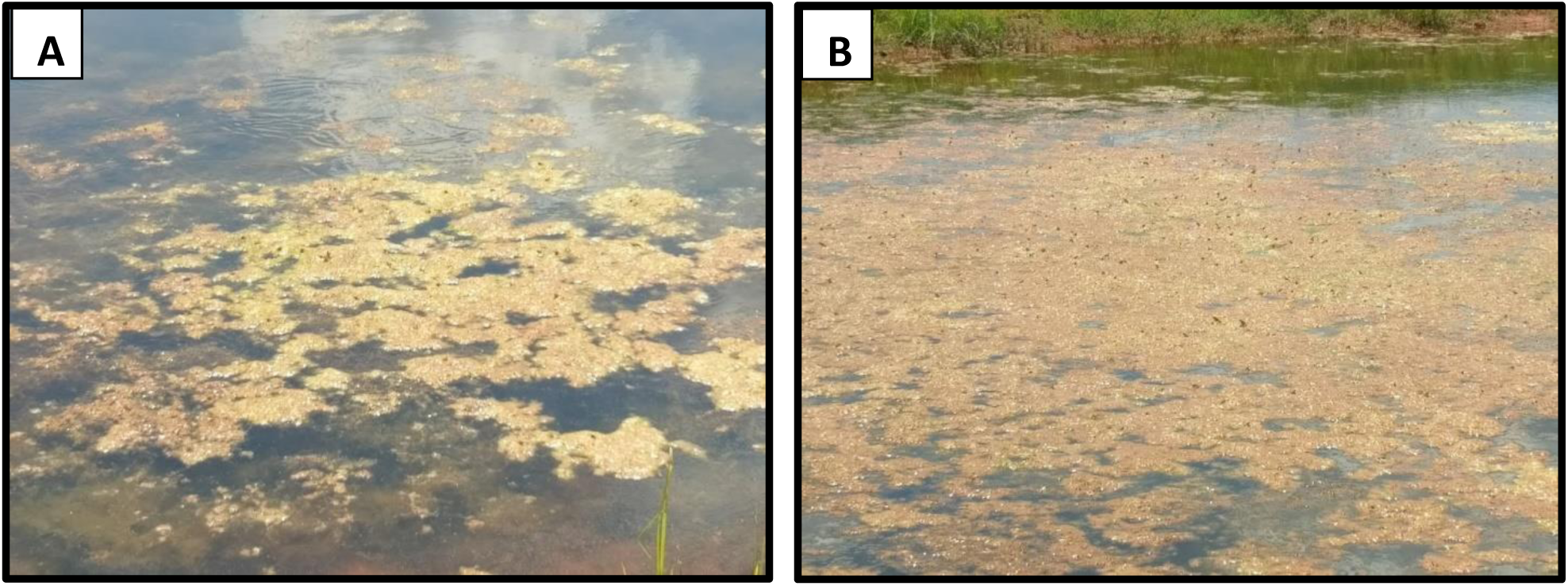
Abundance of Blanchard’s Cricket Frogs (*Acris blanchardi*) in pond microhabitats. A) At least 28 frogs perched on small, thin algal mat (∾3 m long) with submersed *Chara* sp. in Pond A. B) Hundreds of Blanchard’s Cricket Frogs and dozens of juvenile American Bullfrogs (*Lithobates catesbieanus*) utilizing floating algal mats in Pond B. (Video available in supplemental materials).

## Discussion

Previous studies have investigated or commented on the habitat needs of this very adaptable species that can persist in many situations (Martinez-Ortiz 2004). Most accounts suggest these frogs are largely generalized shoreline residents (Conant and Collins 1998). However, I suggest floating mats of vegetation such as *Sphagnum* sp., *Myriophyllum* sp., *Ceratophyllum* sp., duckweed (*Lemna* sp.), *Chara* sp., and filamentous algae (*Spirogyra* sp., *Cladophora* sp.) appear to be important habitat components for Blanchard’s Cricket Frog. At MSNFH, these frogs migrated from one culture pond to another and congregated in areas with this habitat structure, especially during breeding season and during emergence of metamorphs. Here and at many other locations, Duckweed and filamentous algae constitute the primary structural components of these mats. However, the other aforementioned floating plant species appear equally useful.

The slope of the shoreline is important as well. Low-slopes are conducive to breeding choruses and foraging habitat. Habitats along rivers and streams that are steep tend not to serve as habitat. Muddy banks in of themselves do not appear to be a strong predictor of cricket frog occurrence; however, many regions where this species exists are dominated by soil banks and the frogs persist there. Gravel banks tend to be occupied by these frogs. In general, if a habitat has gravel banks, floating mats of vegetation, and a low sloped shore, there is a high probability that it will harbor Blanchard’s Cricket Frog. However, if it has a steep bank and thick cattails or tall grasses along the shore, it is unlikely to be a resident.

Contrary to this study, mud banks were previously identified as primary habitat for this species in West-central Missouri, although proximity to shelter and water played a role (Smith et al. 2003). This region of Missouri is dominated by sandy and muddy creeks and ponds. Some creeks have gravel substrates, but even they commonly have muddy banks. In South Dakota, Blanchard’s Cricket Frog is found in mud and vegetated habitats close to stream edges (Burdick and Swanson 2010). It is possible that differences between our results and laboratory observations reflect the familiarity of the individuals tested and the availability of habitat in West-central Missouri.

A frog that spent its entire life on a muddy bank should be expected to associate with muddy substrates over other types simply because of its familiarity with them (Aitken 1972; Sheldon 1969; Hughes 1997). Availability of habitat may also explain the field observations associating Blanchard’s Cricket Frog with mud banks in Missouri (Smith et al. 2003), South Dakota (Burdick and Swanson 2010), and Illinois (Gray 1984). Most streams and ponds in these specific areas have mud banks. Our study spans these regions, thus providing a wider diversity of habitat types for prediction of abundance and occurrence. This suggests gravel banks as a better predictor of cricket frog occurrence and abundance.

During the summer, cricket frogs will follow floating vegetation mats into open water. At the sphagnum bog in Michigan, we observed Blanchard’s Cricket Frog on sphagnum above the deepest portions of the lake. At Arkansas Post, we observed Blanchard’s Cricket Frog on floating vegetation >50 m from shore. Anecdotal observations at the quaking sphagnum bog in Michigan were at least this from the shoreline. Regan (1973) provided photographs of habitat in Kansas which emulated that of the habitats I tested in this study, I observed at previous locations, and appear present at Glenmere, NY. Removal of thick mats of floating vegetation tends to be the trend in most managed systems, for the convenience of fisherman. The last foothold for Blanchard’s Cricket Frog in New York is a heavily vegetated lake (Glenmere Conservation Coalition, Pers. Comm.). Photographs posted on their site showed ponds with floating sphagnum mats (Available at http://www.glenmere.us, last accessed on 8 June 2009).

In much of the Midwest, where these frogs are declining, the use of herbicides is affecting their persistence. Much attention has targeted direct impacts of these agrichemicals on the reproductive biology and stress response of Blanchard’s Cricket Frog (Reeder et al. 1998; 2005). However, this may be of secondary importance or only a partial explanation of declines. Vegetation mats may serve as important nurseries, based on the large numbers of metamorphs observed in association with them (McCallum et al. 2011). Regan (1973) also noted the use of floating vegetation by Blanchard’s Cricket Frog and commented on its possible importance for tadpoles. Further, floating mats of vegetation allow these frogs to move across ponds and lakes, far from terrestrial and semi-aquatic predators.

The zigzagged escape behavior (McCallum 2011) may be especially effective in aquatic settings when vegetation mats are present. The mats provide refuge, and a multi-colored frog hiding in vegetated mats of duckweed is fairly cryptic. Otherwise, Blanchard’s Cricket Frog should be easy for wading birds and other predators to capture when the complex structure of these mats is not available. Likewise, large frogs like American Bullfrogs may have more difficulty catching a rapidly moving cricket frog while on a vegetation mat compared to in shallow water or on the shore. Cricket frogs are light and appear to jump as effectively on duckweed mats as on land. However, larger frogs and other predators sink into this material and are at an fairly obvious mobility disadvantage. In fact, laboratory findings suggest that if Blanchard’s Cricket Frog is confined in a small space with a predator, it becomes easy prey for predators like bullfrogs (Gray 1978). The evolutionary drive for their jumping behavior depends on the ability to displace and distance itself from an on-looking predator (McCallum 2011). Availability of refuges should facilitate its effectiveness. Vegetation mats may provide unlimited refuge from predators. Males often call from these mats (pers. observ.) and small metamorphs occupying mats are very difficult to find once they bury into the thick floating vegetation (pers. observ.). This is especially critical for early metamorphs that have less developed jumping ability (pers. observ.), because even adult conspecifics will attempt to eat them (McCallum and Trauth 2001).

Many of the chemicals implicated in the declines of Blanchard’s Cricket Frog are herbicides that negatively impact duckweed (Hughes et al. 1988; Hoberg 1993; Solomon et al. 1996; Teodorovic et al. 2012), algae (Fairchild et al. 1998), *Ceratophyllum* sp. (Solomon et al. 1996; Huiyun et al. 2009; Fairchild et al. 1998), and other aquatic plants (Teodorovic et al. 2012; but see Kemp et al. 1985; Jones and Winchell 1984). Therefore, it is quite possible that these herbicides are even more detrimental for cricket frogs when they kill off aquatic vegetation than when they elicit endocrine disruption. Loss of this important habitat component could rapidly lead to extirpation from a locality, especially in light of its short life span (McCallum et al. 2011) simply by exposing them to many small-scale hurdles to survivorship instead of a major catastrophe. First, these chemicals induce physiological stress (Jones et al. 2010), potentially reducing their available energy for predator deterrence and resources for stressor neutralization (McCallum and Trauth 2007). Second, they suppress the development of thick floating mats of vegetation. Third, herbicides disrupt reproduction. I suspect that these three strikes could interact in ways that are more disruptive than any single one unto itself. This may explain why clear identification of a single over-riding factor stimulating declines has been so elusive. It might be prudent for researchers on amphibian declines to not fix their targets on specific individual elements and cast wide nets to find answers. Multiple stressors are almost certainly a more common problem that has thus far been addressed (see Linder et al. 2003). Multiple stressor effects can have additive, facilitative, multiplicative, or even antagonistic interactions with each other (Crain et al. 2008; Folt et al. 1999; Vinebrooke 2004; Newman 2009). So, many small stressors may elicit declines more effectively and more elusively than any single major threat, leading to extirpation or extinction.

Our application of these observations in aquaculture ponds suggests population enrichment is easily accomplished with simple modifications to pond habitat. Management and restoration of this species must include promotion of aquatic vegetation, especially species such as duckweed, *Ceratophyllum* sp. and *Spirogyra* sp. that form dense floating mats. Plant species like *Myriophyllum* sp. that extend up from the substrate and then float along the surface also provide similar habitat. The slope of pond and stream shorelines is also critical. Finally, exclusion of cattails and similar plants from shorelines may assist in the recovery of this frog. Therefore, pond owners should manage small areas of their ponds with these observations in mind.

---

**Appendix 1.** Locations charted for Blanchard’s Cricket Frog (*Acris blanchardi*).

| Location | Abundance*<br>(or number) | Vegetation type<br>where<br>present** | Vegetation type<br>where absent** | Est. Slope from 30 cm<br>above to 30 cm below water<br>edge |  |
| --- | --- | --- | --- | --- | --- |
|  |  |  |  | Present | absent |
| Rice fields, South of Jonesboro, Arkansas, along Highway 1 between N35°43'13.73" W90 41'26.37" and N35°40'18.72" W90 42'15.14". | ++++ | MFV, DW | CTG | < 30° | >30°<br>< 30° |
| Gull Lake, Kellogg Biological Station (KBS), Michigan. (June 1992) | NP |  | SB, LG |  | >30°<br>< 30° |
| Fishing Pier, Red River Research and Education Park, Shreveport, LA (2003 – 2005). N32°26'37.15" W 93°42'5.58" | ++ | LG, DW, MFV | CTG | < 30 | < 30<br>>30 |
| Point 2, Red River Research and Education Park, Shreveport, LA (2003 – 2005). N32°26'41.96" W93°41'51.28" | NP |  | CTG |  | >30<br>< 30 |
| Point 4, Red River Research and Education Park, Shreveport, LA (2003 – 2005). N32°26'36.43" W93°41'25.65" | +++ | LG, MFV | CTG | < 30 | >30<br>< 30 |
| Private lake, Highway 63, Arkansas (Monthly from Oct. 1999 – Jan. 2003.) (N35°52'37.24", W90°47'26.06") | ++++ | MFV, MF, DW | CTG | < 30° |  |
| Quaking Bog, vic. Of KBS. (June 1992) | +++ | SFM, WL | CTG | --*** | -- |
| Betty Spring, vic. Lake Frierson, Jonesboro, AR. (Summer 2001). N35°58'20.59" W90°44'06.70" | +++++ | MFV, SV, LG |  | < 30° |  |
| Private pond, vic. Normal, IL. May 1992. | ++ | MFV, MF, DW |  | >30 |  |
| Sand Pond, Savanna Army Depot, Jo Daviess County, IL (Summer 1994). N42°13'55.72" W90°19'13.16" (large bullfrog tadpole population) | NP |  | CTG, RS, SB, MF, DW |  | all |
| Creek, Savanna Army Depot, Jo Daviess County, IL (Summer 1994). N42°15'48.25 W90°22'08.57" (Large populations *L. pipiens*, *L. Palustris*). | NP |  | TG, MF |  | >30° |
| Agricultural Ditch north of Bono, AR, crossing Highway 63 (Monthly, Oct. 1999 – Jan. 2003). N35°54'58.31" W90°48'18.70" | +++ | MFV, DW, LG | MF, LG | < 30° | >30° |
| Agricultural Ditch south of Jonesboro, AR, along Highway 1, N35°40'55.50" W90°42'15.93" | ++++ | DW, CTG | MFV, CTG |  | >30° |
| Campsite, Savanna Army Depot, Jo Davies County, IL (Summer 1994) N42 13'10.10" W90 20'38.55" | NP | LG, CTG |  |  | all |
| Salado Creek, Arkansas (Summer 2000) N35°39'36.17", 91°34'06.15" | 1 | MF |  |  | >30° |
| Dawt Mill Resort, Black River, Missouri. West bank, north and south of bridge. (1999 – 2003) N36°36'35.92" W92°16'43.05" | +++++ | GB, GBW, SV |  | < 30<br>>30 |  |
| Dawt Mill Resort, Black River, Missouri, north and south of bridge (1999 – 2003). N36°36'32.98" W92°16'42.79" | + | RG, RGV |  |  | >30 |
| Giles Creek, Collinsville, IL (1979 – 2003), | + | MFV, LG | CTG, MF |  |  |
| N38°42'42.36" W89°56'51.84" |  |  |  |  |  |
| Rex Dr. Lake, Collinsville, IL (1979 – 1984)<br>N38°42'48.45" W89°56'47.76" | +++ | DW, CTG | CTG, MF | >30 | >30 |
| Lakeview Acres I, Collinsville, IL (1979 – 1999),<br>N38°42'41.24" W89°56'58.34" | NP |  | MF, RS, LG |  | >30 |
| Lakeview Acres II, Collinsville, IL (1983 – 1986),<br>N38°42'47.85" W89°56'59.51" | NP |  | RS, LG |  | >30 |
| Pasture Creek, Maryville IL (1983 – 1984)<br>N38°43'04.22" W89°56'21.88" | ++ | LG | MF, CTG | < 30 | >30 |
| Wooded Creek, Maryville, IL (1983 – 1984)<br>N38°43'03.81" W89°56'27.69" | +++ | GBW, GB,<br>MFV |  | < 30 |  |
| Wooded Creek, Maryville, IL (1983 – 1984)<br>N38°42'40.69" W89°56'29.29" | NP |  | MF |  | >30 |
| Holden Lake, Holden, Missouri (2012 -2016)<br>N38°45'52.07" W94°02'22.03", N38°45'26.94"<br>W94°01'54.99", N38°45'16.18" W94°02'05.63" | + | MF, LG, | RS, WL, CTG | < 30 | >30 |
| Spillway, Lake Wright Patmann, Texarkana, Texas<br>(2003 – 2011) N33°18'21.02" W94°09'24.65" | 1 | RS | RS, MF, CTG | >30 | >30 and <30 |
| Waterfowl observation post, Lake Wright Patmann,<br>Texarkana, Texas (2005 – 2011) N33°19'42.49"<br>W90°09'41.81" | ++++ | MFV, DW | CTG, RS | < 30 | >30<br>< 30 |
| Jane's Creek (1999 – 2003) northeast of Ravenden<br>Springs, AR at<br>N36°23'59.24" W91°15'49.71" | +++++ | MFV, GBW,<br>SV, GB | GB, CTG, | < 30 | >30 |
| Livestock pond at home (2012 – 2016)<br>N38°47'13.06" W94°02'02.89" | +++++ | DW MFV MF<br>LG | CTG | < 30 | >30 |
| Home reservoir (2012 – 2016)<br>N38°47'16.36" W94°02'10.69" | ++ | DW MF MFV<br>LG | CTG LG MF | < 30 | >30 |
| Wooded Creek (2012 – 2016)<br>N38°47'41.52" W94°01'42.85" | +++ | GB, GBW,<br>MFV | GBW, MF | < 30 | >30 |
| Silver Lake Park, Highland, Illinois (1980 – 1987)<br>N38°46'01.49" W89°41'38.99" | + | CTG, LG, SV,<br>DW | CTG, RS, LG | >30 | >30 |
| Wooded pond, Southern Illinois University at<br>Edwardsville campus (1995-1999) N38°47'32.36"<br>W89°59'16.64" | NP |  | MF |  | >30 |
| Langston University Campus ponds (20016 – 2017) | ++ | DW, MFV, LG<br>SV | CTG, RS, MF<br>LG | < 30 >30 | >30 |
| Pasture spring-pond along US 412 north of Black<br>Rock, AR. (1999 – 2002). N36°07'19.05"<br>W91°08'55.15" | ++++ | MFV, LG, MF<br>DW | CTG | < 30 | >30 < 30 |
| Maryville Fishing Club Lake, Maryville, Illinois<br>(1983 – 2002)<br>N38°43'41.33" W89°56'58.70" | ++++ 1983<br>+ 2002 | MFV, DW, LG<br>MF | RS, MF, CTG | < 30 | >30 |
| White Oak Park Lake, Normal, Illinois (1989 – 1992)<br>N40 29'38.43" W89 00'53.25" | +++++ | MFV, LG |  | < 30 |  |

\*Abundance: NP = not present, + = rare, ++ =not common , +++ = common, ++++ = very abundant
\*\*Key to vegetation types:
GB = gravel bed without vegetation
GBW = gravel bed with vegetation
RG = bedrock with some gravel
RGV = bedrock with some gravel and some taller vegetation.
LG = lawn grass to edge of water
CTG = thick bed of Cattails or tall grasses in water and along shoreline.
RS = rock shoreline, rubble for erosion control
SB = sand bottom, no shallow water vegetation, grass may be present on shoreline
MF = mud flat without aquatic vegetation, grasses or rushes were present within 30 cm of shore.
MFV = mud flat with aquatic or emergent vegetation including small rushes and grasses extending into the water.
SFM = sphagnum floating mats
DW = floating duckweed, algal mats, or similar vegetation.

**Appendix 2.** Multiple linear regression results for each of the five best models associating habitat observations with abundance of Blanchard,s Cricket Frog (*Acris blanchardi*). (GBW = gravel banks without vegetation, SFM = Sphagnum mats, DW = duckweed-algal mats, SV = thick submerged floating mats of *Ceratophyllum* sp. or *Myriophyllum* sp., slope = shoreline slope > 30°).

|  |  |  |  |  |  |  |
| --- | --- | --- | --- | --- | --- | --- |
| **Model 1** | Abundance = 1.82 + 1.43 GBW + 2.80 SFM + 1.23 DW + 1.50 SV – 1.62 slope.<br>R<sup>2</sup> = 0.573. Durbin-Watson = 1.71. No evidence of lack of fit (P > 0.1). |  |  |  |  |  |
|  | **Predictor** | **Coef** | **SE Coef** | **T** | **P** | **VIF** |
|  | Constant | 1.8213 | 0.3590 | 5.07 | < 0.001 |  |
|  | GBW | 1.4262 | 0.6424 | 2.22 | 0.031 | 1.3 |
|  | SFM | 2.8020 | 1.2310 | 2.28 | 0.027 | 1.0 |
|  | DW | 1.2286 | 0.3997 | 3.07 | 0.003 | 1.3 |
|  | SV | 1.5044 | 0.5701 | 2.64 | 0.011 | 1.2 |
|  | Slope | -1.6232 | 0.3711 | -4.37 | < -0.001 | 1.2 |
|  | **Analysis of Variance** |  |  |  |  |  |
|  | **Source** | **Df** | **SS** | **MS** | **F** | **P** |
|  | Regression | 5 | 104.550 | 20.910 | 14.23 | < 0.001 |
|  | Residual error | 53 | 77.857 | 1.469 |  |  |
|  | Lack of fit | 5 | 24.901 | 4.980 | 4.51 | 0.002 |
|  | Pure error | 48 | 52.956 | 1.103 |  |  |
|  | Total | 58 | 182.407 |  |  |  |
| **Model 2** | Abundance = 1.82 + 143 GB + 2.80 SFM + 1.23 DW + 1.50 SV – 1.62 slope<br>R<sup>2</sup> =0.573. Durbin-Watson = 1.93. No evidence of lack of fit (P > 0.1) |  |  |  |  |  |
|  | **Predictor** | **Coef** | **SE Coef** | **T** | **P** | **VIF** |
|  | Constatant | 1.8213 | 0.3590 | 5.07 | < 0.001 |  |
|  | GB | 1.4262 | 0.6424 | 2.22 | 0.031 | 1.3 |
|  | SFM | 2.8020 | 1.2310 | 2.28 | 0.027 | 1.0 |
|  | DW | 1.2286 | 0.3997 | 3.07 | 0.003 | 1.3 |
|  | SV | 1.5044 | 0.5701 | 2.64 | 0.011 | 1.2 |
|  | slope | -1.6232 | 0.3711 | -4.37 | < -0.001 | 1.2 |
|  | **Analysis of Variance** |  |  |  |  |  |
|  | **Source** | **Df** | **SS** | **MS** | **F** | **P** |
|  | Regression | 5 | 104.550 | 20.910 | 14.23 | < 0.001 |
|  | Residual error | 53 | 77.857 | 1.469 |  |  |
|  | Lack of fit | 5 | 24.901 | 4.980 | 4.51 | 0.002 |
|  | Pure error | 48 | 52.956 | 1.103 |  |  |
|  | Total | 58 | 182.407 |  |  |  |
| **Model 3** | Abundance = 1.24 + 1.32 GBW + 0.912 MFV + 2.81 SFM + 0.915 DW + 1.29 SV – 1.05 slope<br>R<sup>2</sup> = 0.593. Durbin-Watson = 2.02<br>Possible curvature in variable DW ( P = 0.011)<br>Possible interactions with variable DW (P = 0.012)<br>Possible interactions with variable SV (P = 0.026)<br>Possible curvature in variable “slope” (P = 0.061)<br>Possible lack of fit at outer x-values (P = 0.029)<br>Overall lack of fit test is significant (P = 0.011) |  |  |  |  |  |
|  | **Predictor** | **Coef** | **SE Coef** | **T** | **P** | **VIF** |
|  | Constant | 1.2413 | 0.5077 | 2.45 | 0.018 |  |
|  | GB | 1.3169 | 0.6370 | 2.07 | 0.044 | 1.3 |
|  | MFV | 0.9117 | 0.5721 | 1.59 | 0.117 | 3.1 |
|  | SFM | 2.8120 | 1.2140 | 2.32 | 0.024 | 1.0 |
|  | DW | 0.9145 | 0.4406 | 2.08 | 0.043 | 1.6 |
|  | SV | 1.2900 | 0.5779 | 2.23 | 0.030 | 1.3 |
|  | slope | -1.0529 | 0.5117 | -2.06 | 0.045 | 2.4 |
|  | **Analysis of Variance** |  |  |  |  |  |
|  | **Source** | **Df** | **SS** | **MS** | **F** | **P** |
|  | Regression | 6 | 108.1750 | 18.029 | 12.63 | < 0.001 |
|  | Residual error | 52 | 74.2320 | 1.428 |  |  |
|  | Lack of fit | 8 | 33.2860 | 4.161 | 4.47 | 0.001 |
|  | Pure error | 44 | 40.9450 | 0.931 |  |  |
|  | Total | 58 | 182.4070 |  |  |  |
| **Model 4** | Abundance = 1.24 + 1.32 GB + 0.912 MFV + 2.81 SFM + 0.915 DW + 1.29 SV – 1.05 slope<br>R<sup>2</sup> = 0.593. Durbin-Watson = 2.02<br>Possible interactions with variable DW (P = 0.001) |  |  |  |  |  |

Possible lack of fit at outer x-values (P = 0.022) Overall lack of fit test is significant (P = 0.001)
| Predictor | Coef | SE Coef | T | P | VIF |
| --- | --- | --- | --- | --- | --- |
| Constant | 1.2413 | 0.5077 | 2.45 | 0.018 |  |
| GB | 1.3169 | 0.6370 | 2.07 | 0.044 | 1.3 |
| MFV | 0.9117 | 0.5721 | 1.59 | 0.117 | 3.1 |
| SFM | 2.8120 | 1.2140 | 2.32 | 0.024 | 1.0 |
| DW | 0.9145 | 0.4406 | 2.08 | 0.043 | 1.6 |
| SV | 1.2900 | 0.5779 | 2.23 | 0.030 | 1.3 |
| slope | -1.0529 | 0.5117 | -2.06 | 0.045 | 2.4 |

Analysis of Variance
| Source | Df | SS | MS | F | P |
| --- | --- | --- | --- | --- | --- |
| Regression | 6 | 108.1750 | 18.029 | 12.63 | < 0.001 |
| Residual error | 52 | 74.2320 | 1.428 |  |  |
| Lack of fit | 8 | 33.2860 | 4.161 | 4.47 | 0.001 |
| Pure error | 44 | 40.9450 | 0.931 |  |  |
| Total | 58 | 182.4070 |  |  |  |

| Predictor | Coef | SE Coef | T | P | VIF |
| --- | --- | --- | --- | --- | --- |
| Constant | 1.4071 | 0.5238 | 2.69 | 0.010 |  |
| GBW | 1.3065 | 0.6342 | 2.06 | 0.045 | 1.3 |
| MF | -0.3962 | 0.3285 | -1.21 | 0.233 | 1.1 |
| MFV | 0.8438 | 0.5724 | 1.47 | 0.147 | 3.1 |
| SFM | 2.6310 | 1.2180 | 2.16 | 0.035 | 1.0 |
| DW | 0.9993 | 0.4443 | 2.25 | 0.029 | 1.7 |
| SV | 1.1869 | 0.5817 | 2.04 | 0.046 | 1.3 |
| Slope | -1.0386 | 0.5097 | -2.04 | 0.047 | 2.4 |

Analysis of Variance
| Source | Df | SS | MS | F | P |
| --- | --- | --- | --- | --- | --- |
| Regression | 7 | 110.234 | 15.748 | 11.13 | < 0.001 |
| Residual error | 51 | 72.173 | 1.415 |  |  |
| Lack of fit | 12 | 40.916 | 3.410 | 4.25 | < 0.001 |
| Pure error | 39 | 31.256 | 0.801 |  |  |
| Total | 58 | 182.407 |  |  |  |

**Appendix 3.**
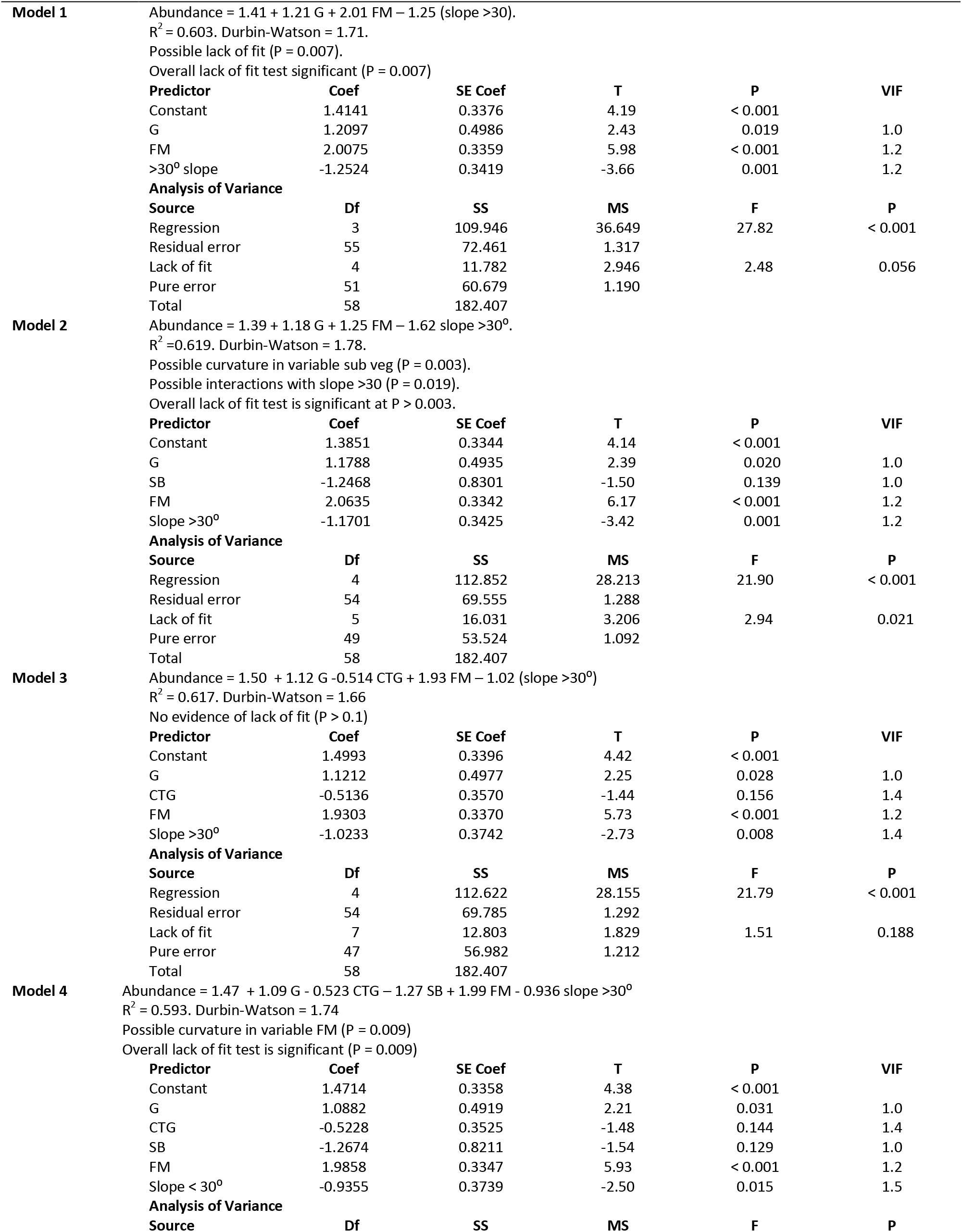
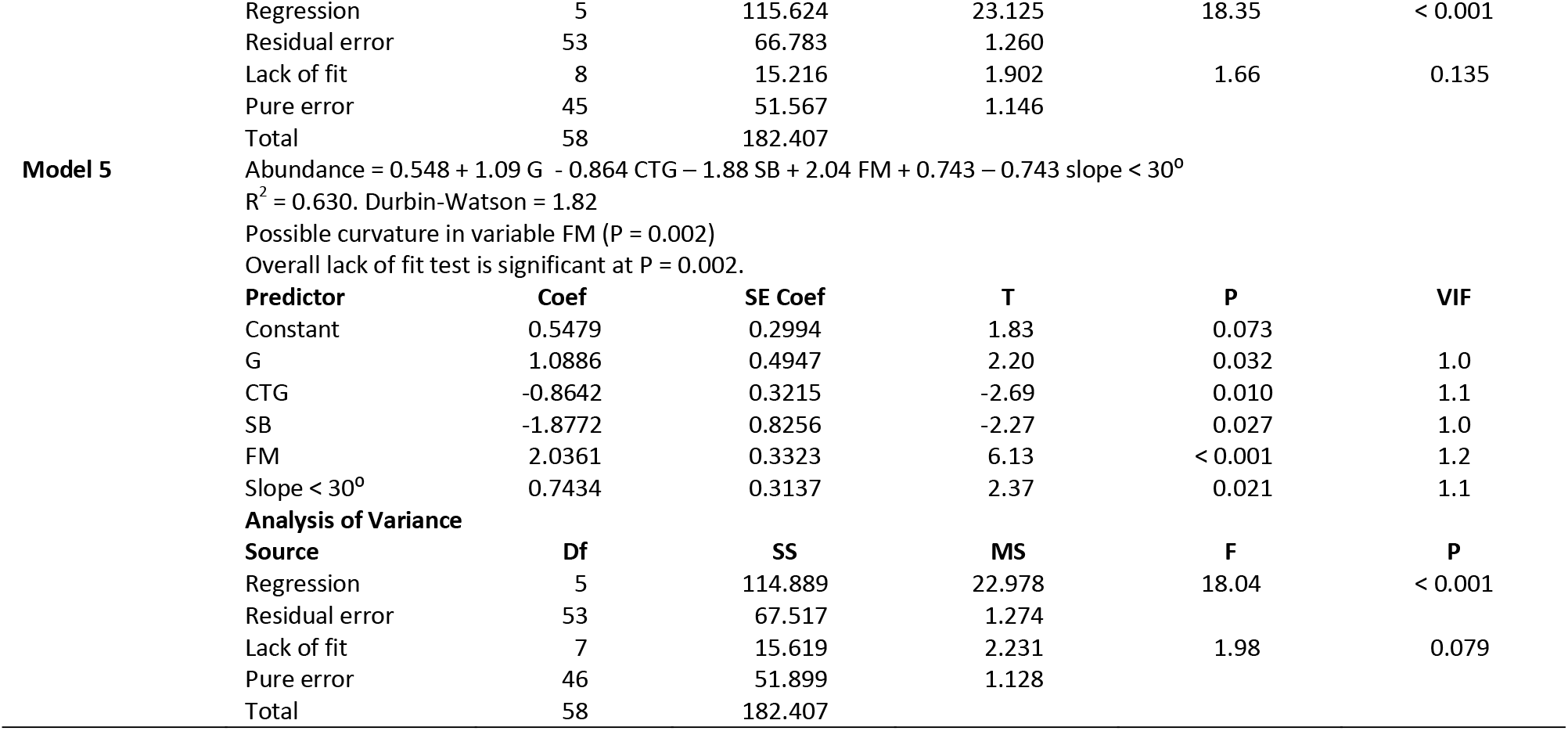
Multiple linear regression results for each of the five best models associating consolidated habitat observations with abundance of Blanchard,s Cricket Frog (*Acris blanchardi*). (G = gravel banks; SF = floating vegetation, slope = shoreline slope > 30° or < 30°).

## SUPPLEMENTAL MATERIALS

Videos of Blanchard’s Cricket Frog (*Acris blanchardi*) abundance along the shorelines of these ponds to demonstrate the robust population size after application of field observations to mock restoration activities. Abundance is clearly more robust in Pond B. (Available at: TBA by HCB editorial staff).

VIDEO0001.MP4 = northwestern corner of pond B, VIDEO0002.MP4 = northern shore of pond B,

VIDEO0004.MP4 = northern shore of pond A, VIDEO0005.MP4 = southwestern corner and south shore of pond B,

VIDEO0005.MP4 = vegetation mat and adjacent west shoreline pond B, VIDEO0010.MP4 = west shore of pond A,

VIDEO0011.MP4 = southern shore of pond A, VIDEO0012.MP4 = algal mats in pond B.

